# Hydraulic Conductivity of The Hemp Stems Under Water Stress

**DOI:** 10.1101/2023.06.09.544341

**Authors:** Mehmet Yuksel, Hadi A. Al-agele, Lloyd Nackley, Chad W. Higgins

## Abstract

The hydraulic conductivity of hemp stems under water-stress conditions was investigated to assess the impact on water transport from the root to the leaves. Water-stress conditions induce embolism (cavitation) in xylem channels, thereby affecting water flow. The percentage loss of water transfer ability within the xylem channels can be represented by a ‘vulnerability curve’ (VC). This study utilized an air injection technique to induce embolisms in the stem and measured the subsequent changes in hydraulic conductivity using a specialized measurement apparatus. The results revealed that the shape of the vulnerability curve for hemp was influenced by the xylem area, which is an atypical finding with no prior evidence in other plant species. The statistical analysis confirmed the significance of this effect (p-value=0.003 at 95% confidence intervals). Consequently, a non-traditional mathematical equation (simple power law) was developed to describe the relationship between pressure and xylem area. Furthermore, our findings demonstrated that hemp stems are more responsive to water stress compared to other plant species documented in previous literature. The minimum pressure (0.15 MPa) at which initial cavitation is observed and the maximum pressure (∼2.4 MPa) at which all stem conductance is lost were among the lowest reported values, indicating that hemp growth may be significantly affected by deficit irrigation strategies. Thus, careful monitoring of the irrigation schedule is crucial, particularly for younger stems with smaller xylem cross-sectional areas. These insights into the hydraulic behavior of hemp can contribute to the development of improved irrigation strategies in agriculture, ultimately enhancing water-use efficiency and optimizing crop production.

**Highlight:**

1. The hydraulic conductivity of hemp stems under water-stress conditions was investigated to assess the impact on water transport from the root to the leaves.
2. The results revealed that the shape of the vulnerability curve for hemp was influenced by the xylem area, which is an atypical finding with no prior evidence in other plant species.
3. Our findings demonstrated that hemp stems are more responsive to water stress compared to other plant species documented in previous literature.
4. The minimum pressure (0.15 MPa) at which initial cavitation is observed and the maximum pressure (∼2.4 MPa) at which all stem conductance is lost were among the lowest reported values, indicating that hemp growth may be significantly affected by deficit irrigation strategies.
5. Thus, careful monitoring of the irrigation schedule is crucial, particularly for younger stems with smaller xylem cross-sectional areas. These insights into the hydraulic behavior of hemp can contribute to the development of improved irrigation strategies in agriculture, ultimately enhancing water-use efficiency and optimizing crop production.

## Introduction

Population growth and climate change are major contributors to global water stress, and it is anticipated that by 2050, an additional 1.8 billion people will live in the water stresses area (Schlosser et al., 2014). Concerns of water scarcity are prevalent and fresh water makes up only around 2.5 percent of the world’s total water (Oki & Kanae, 2006). Considering these limitations, efficient and effective water management across all its major uses is critical.

Irrigation is the largest consumer of water globally. It accounts for approximately 70 percent of the water consumption in most of the regions of the world (Khokhar, 2017) with higher percentages (∼80%) in arid and water-stressed, dry and semi-arid regions (Fereres & Soriano, 2007). Efficient water-use in agriculture; therefore, becomes a major research focus.

Agricultural water management research seeks to enhance water usage efficiency (Jones, 1990) through improved technologies or management practices. More recently, these efforts have extended to the water-energy-food nexus as it is obvious that sustainable food production cannot be sacrificed for sustainable water management. That is, efforts have sought to achieve both sustainable water management and sustainable agriculture production (Al-agele et al., 2022; Al-agele et al., 2020; AL-agele et al., 2021a, 2021b, 2021c; Irmak et al., 2011; Qin et al., 2011; Smathers et al., 1995). One such strategy is deficit irrigation (DI) where irrigation water is applied at a rate below what is necessary for evapotranspiration (ET). Water conserved can be redirected for alternative uses (Fereres & Soriano, 2007). One pitfall of DI is that it can induce water stress in plants, and effective implementation of a DI strategy requires managing plant water stress.

Plants are frequently exposed to biotic and abiotic stresses in nature and water stress is one of the most substantial hindrances to plant development and productivity (Anjum et al., 2011). Therefore, water stress, and it’s management, is a major issue in agriculture (not just for DI), and the capacity for plants to endure such stress is extremely important economically (Shao et al., 2008). Water stress triggers negative physiological effects in plants that are both morphological effects and biochemical (Anjum et al., 2011). The initial response to water stress is stomatal closure, which decreases photosynthesis (Anjum et al., 2011; Osakabe et al., 2014). Longer exposures to stress can alter the number of leaves per plant, leaf size, leaf life, and reduce stem water conductance (Anjum et al., 2011), and dramatically decrease yields (Anjum et al., 2011; Onder et al., 2005; Tesfamariam et al., 2010).

The focus of this paper is the loss of stem water conductance in the presence of water stress. In plants, water evaporates from leaves and tension is created by that evaporation. This tension, in turn, causes negative water potential and the water is pulled from roots to leaves through the stem by liquid water’s cohesive strength (Cochard et al., 2013; Sheridan & Nackley, 2021). This hydraulic pathway is interrupted when the cohesive strength of the water’s tension is broken with an embolism (vapor pocket within the stem’s xylem). The decrease in stem water conductance is due to the formation of water vapor and air intrusions in xylem channels called embolism (Hargrave et al., 1994; Sperry & Tyree, 1990; Vilagrosa et al., 2020) and the drop in xylem conductivity caused by vessel embolism can cause a reduction in water flow in xylem.

Prior studies have shown that the water flow rate plants through stems changes when they are under water stress (Lovisolo & Schubert, 1998) because an increases in plant water stress caused a decrease in stem water potential (McCutchan & Shackel, 1992). As a consequence, stem water conductance reduces (Bucci et al., 2003; Thomas & Eamus, 1999). Hydraulic characteristics of the stem xylem have been measured to provide fundamental information of the plant’s capability to procurement water for photosynthetic and growing textures (Brodribb, 2009; Holbrook & Zwieniecki, 2005; Tyree & Ewers, 1991). Lovisolo & Schubert found that the plants which were irrigated have 35% higher stomal conductance than water stressed plants (Lovisolo & Schubert, 1998). Also, the shoot hydraulic conductivity reduced by approximately 42-72% in water stressed conditions. De Silva et al., investigated the effects of water stress and heavy metal stress on hydraulic conductivity in red maple (Acer rubrum L.) (De Silva et al., 2012). They found the xylem-specific stem conductivity decreased by 40-50% in both stress conditions. Gleason et al., found a 50%-80% decrease in whole-plant hydraulic conductivity in Zea Mays when it was exposed to water-stressed conditions (Gleason et al., 2017).

*Cannabis sativa* (cannabis) was one of the first plants grown by human-being (Zuardi, 2005). From a historical perspective, since the Neolithic era, cannabis has been discovered in China approximately 6000 years ago (Li, 1974). Today, the cannabis plant polarizes societies with social and political prejudices (Afrin et al., 2020). Little information is available within the literature on the stem water conductance of hemp underwater stress. Tang et al., investigated water and nitrogen use efficiencies of hemp and showed that plants under water stress had less stem, root, and green leaf biomass (Tang et al., 2018). García-Tejero et al., performed a deficit irrigation trial with full irrigation and a 20% deficit. Their results showed yield reduction with this deficit (García-Tejero et al., 2014). Gill et al., cultivated hemp under water stress and demonstrated that hemp was able to survive at poor availability of water in the soil (Gill et al., 2022). However, the biomass production, root dry weight, shoot dry weight, height of hemp plant and total leaf area per plant reduce.

This research aims to measure the change in hemp stem water conductivity in response to imposed stress, specifically through the forced creation of embolisms. A simple mathematical expression will be fitted to these measurements, and the consequences of these findings will be explored.

## 1. Material and Method

### 1.1 Plant Breading

In this study, we investigate the changes in hydraulic conductivity of hemp stems under water stress conditions. The hemp plants were cultivated in the greenhouses at North Willamette Research and Extension Center in Aurora, OR 97002. The plants were grown in small pots with volumes of 4” (0.5 L) and 7” (4.5 L), respectively. To maintain their water requirements, the plants were irrigated twice a day using an automatic sprinkler system at 8:00 AM and 3:00 PM. The duration of irrigation was adjusted between 2-5 minutes based on prevailing weather conditions. Additionally, a weekly application of 200 ppm N fertilizer was administered using a hand watering-can.

The hemp plants used in this study had varying ages, ranging from 2 months to 6 months old. The overall heights of the plants ranged between 48 cm and 92 cm, while the main stem diameters varied from 0.37 cm to 0.82 cm. To calculate the xylem area, pith diameters were also measured, which ranged between 0.08 cm and 0.27 cm. The plants were transported alive to the NEWAg Lab in Weniger Hall at Oregon State University, where they were harvested. Data collection and measurements commenced immediately after harvesting. All plant materials were harvested and measured during the period of July to August 2022.

A general overview of the procedural steps is a follows: a healthy stem is chosen; a leaf cluster attached to the chosen stem is prepared and removed from the chosen stem; the water potential of the now removed leaf cluster is measured with a PMS 600 pressure chamber; the chosen stem is sacrificed and prepared for water conductivity measurements; the ‘natural’ stem water conductivity is measured; all embolisms are removed from the stem with a vacuum chamber; the maximum conductance is measured; the stem sample is then exposed to increasing levels of pressure within the PMS 600 pressure chamber (to simulate a stress condition) and the stem water conductance measurements are repeated until no water passes through the stem.

The flowchart shows all the processes of the study.

Stems with visible damage, pest prevalence or other disorders that may induce stress along the entire stem (even outside the proposed measurement section) are excluded. The leaves on a selected stem are then covered by aluminum foil to prepare for the stem water potential measurement. Sample stems with leaves wrapped, are left to acclimate for one hour after which the leaf groups are removed with a sharp razor blade. The timing of the leaf sample harvest was set to midday for all collected samples (Bucci et al., 2003; Holtzman et al., 2021). These leaf clusters are sent to a PMS instrument’s model 600 pressure chamber (Scholander et al., 1965) to measure the leaf water potential (leaf Ψ) and stem water potential (stem Ψ) (Choné et al., 2001).

An approximately 15 cm sample was then taken from the harvested stem. All stem samples were cut neatly with a sharp razor whilst the stem sections were submerged in water. This reduces the chance that the xylem is contaminated with extra air bubbles (Sperry & Tyree, 1990; Sperry, Donnelly, & Tyree, 1988). All harvested stems were defoliated, and wood glue was applied to areas of the stem’s section containing a removed branch. These same areas were then wrapped in parafilm. This was done to prevent water dripping or leaking laterally during hydraulic conductivity measurement. Finally, the flow direction, from larger cross-sectional area to the thinner cross-sectional area, was marked and was maintained consistently throughout all measurements.

The water flow rate through the stem in the measurement apparatus can become too slow or near zero if the stem length is too long. This is undesirable as it creates a time impediment to the experiments, and previous studies have shown that stem length did not impact the conductivity measurements (Pérez-Harguindeguy et al., 2013)(Choat et al., 2015; Cochard et al., 2010). Nevertheless, length of stem section was adjusted accordingly with the ‘vessel-length procedure’ (Pérez-Harguindeguy et al., 2013). Here, ‘vessel-length’ refers to the length of a single vessel in the xylem of the plant stem. This procedure is described briefly below and shown visually in Figure 3.

**Figure 1.**
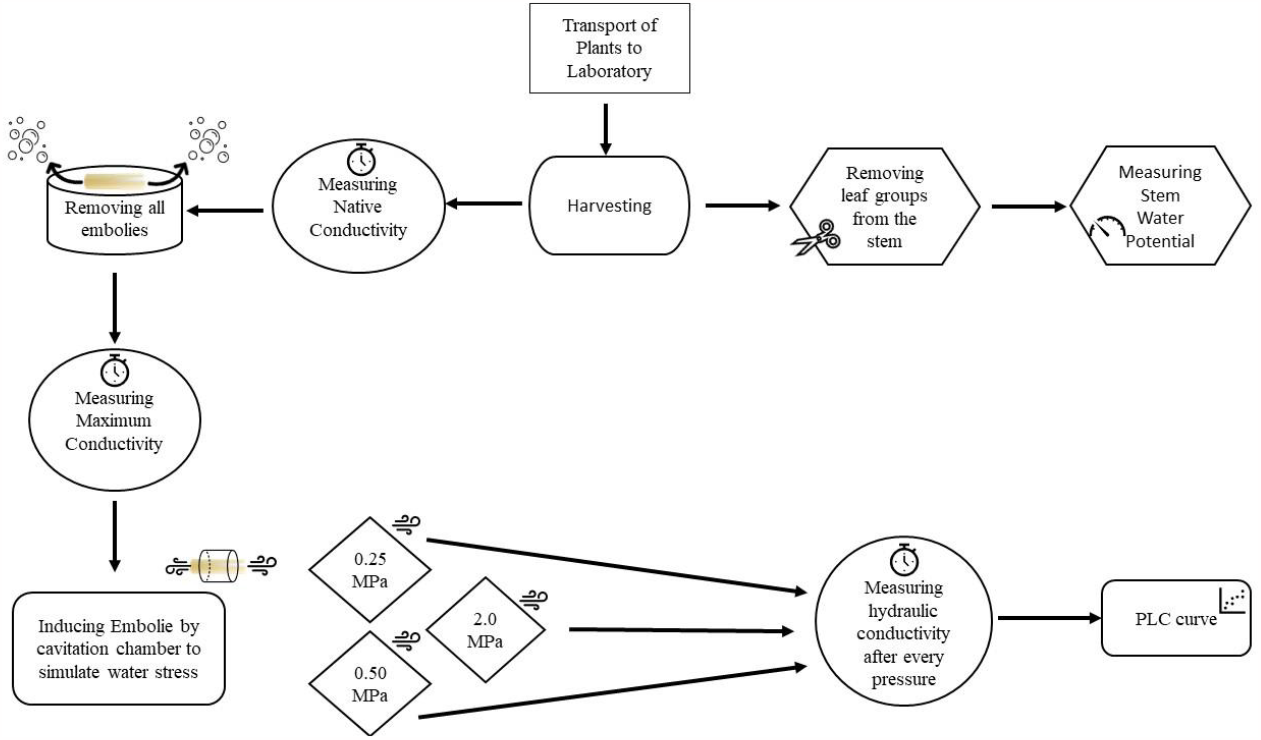
All the processes of the hydraulic conductivity project

**Figure 2.**
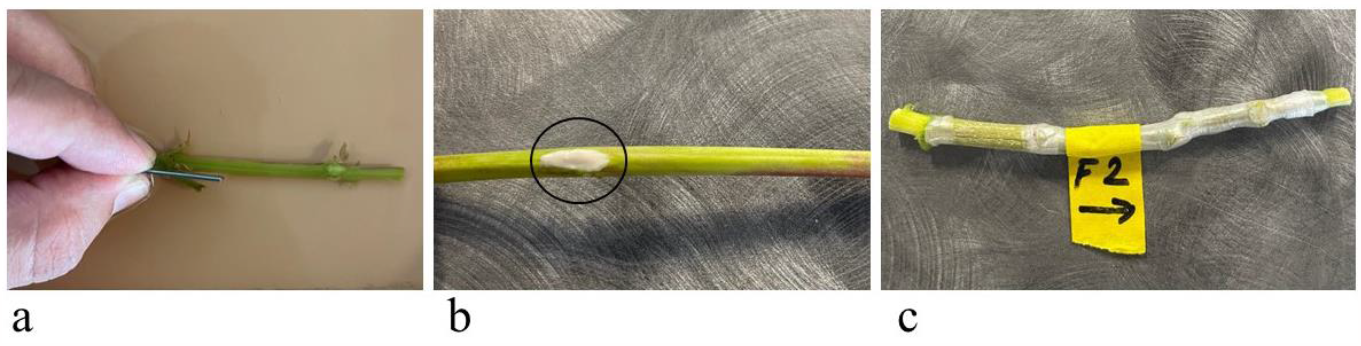
a) removing brunches from the sample, b) applying wood glue, c) marking flow direction and wrapping by parafilm

**Figure 3.**
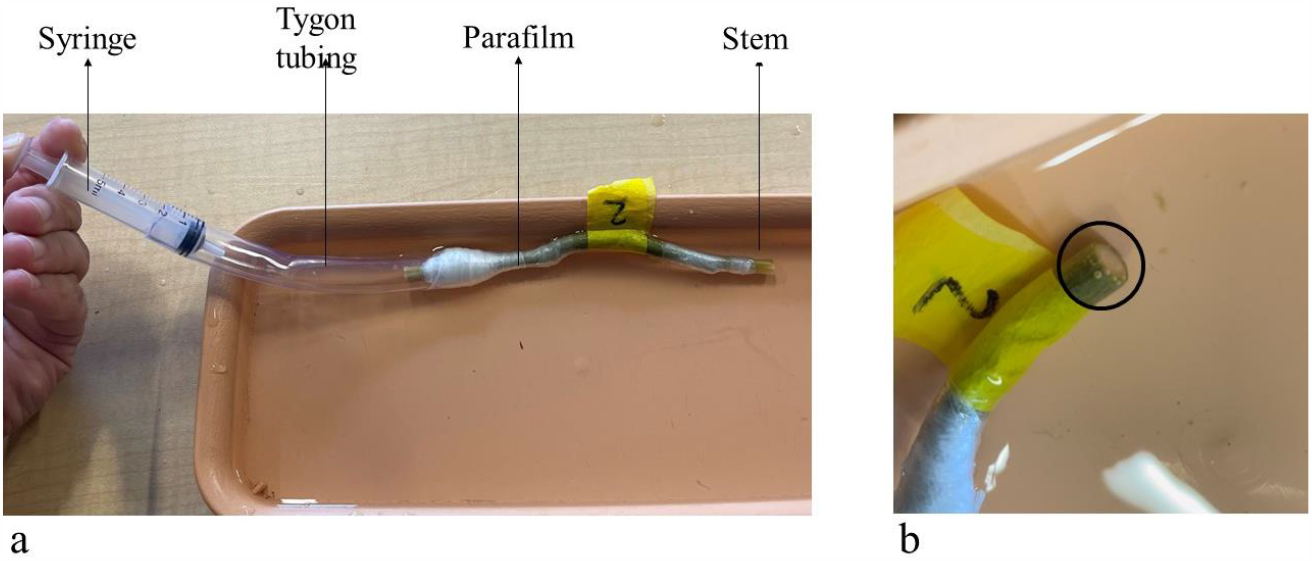
a) applying the air by pressure, and b) visible air bubbles.

A syringe is attached to one end of the submerged sample and compressed. If bubbles could be seen at the opposite end of the stem, the stem is shorter than the maximum xylem vessel length (Pérez-Harguindeguy et al., 2013). Our aim was to find the threshold-length at which bubbles first appear. This was achieved by starting with a long stem and repeating this test after shortening the stem by ∼2mm until bubbles were visible (Figure 3).

The stem sample must be hydraulically connected to the measurement apparatus after the measurement apparatus is prepared and its elements are calibrated. A proper fit was found by 1) adjusting the size of the tygon tubing and 2) wrapping stem ends in laboratory parafilm to fit the tygon tubing. Air bubbles can also appear in the connection. To prevent air bubbles, connections are made while water flows through the system. See Figure 4a for a well-executed connection and Figure 4b for a photo of a connection with an air bubble that must be redone.

**Figure 4.**
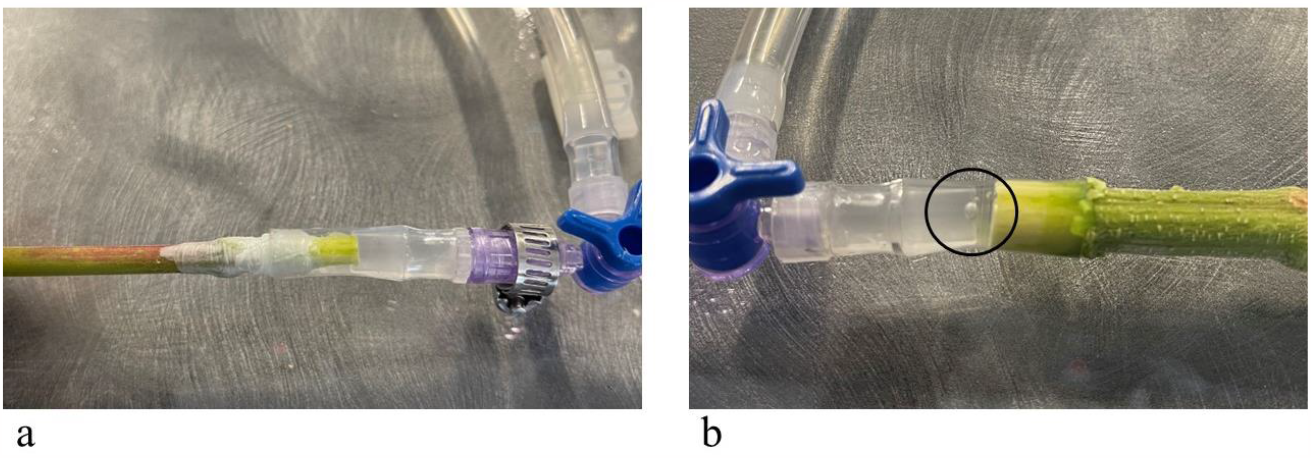
a) correct connection to the system and b) wrong connection because of air bubble.

### 1.2 Hydraulic Conductivity Measurement Apparatus

A gravity hydraulic flow apparatus was used to measure the stem hydraulic conductivity. Minor modifications of Sperry’s original device were made: we used a manual balance instead of a computer controlled balance, and we added standing pickets to further visualize pressure head (Sperry et al., 1988).

The system contains an elevated reservoir of water that feeds De-Ionized (DI) water to a test section of a stem. The elevation difference between the water reservoir and the stem connection section creates a pressure head which drives water flow through the stem test section. The water passes through the plant stem exits to a precision balance (see Figure 5). The rate of water accumulation on the balance is used to compute the flow rate, and the flow rate and applied pressure head is utilized to calculate hydraulic conductivity.

**Figure 5.**
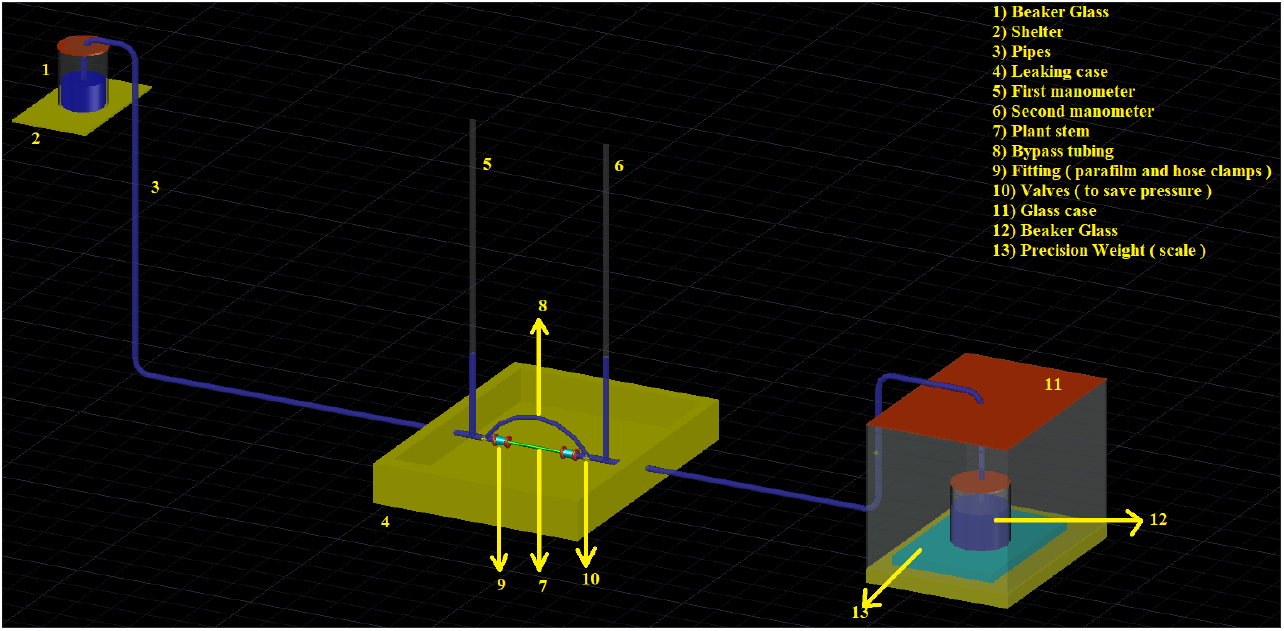
Illustrates water flow system to measure hydraulic conductivity of the stems.

An illustration of the measurement apparatus is presented in Figure 5.

A two decimal per gram (1.00 gram) precision scale was utilized. Note that the ultimate precision in the hydraulic conductivity measurements is related to the precision in the scale utilized. The calibration of the scale was checked prior to each measurement with standard weights of 1, 2, 3, 5, and 10 g respectively. The water temperature was measured for every associated stem conductivity measurement, and the water density was calculated based on the temperature. This density is used in equation 1.

The system is fully driven by gravity, and the reservoir height was chosen to supply ∼8 kPa (pressure head) for all samples. This pressure was chosen because it is lower than a pressure that would remove emboli (Choat et al., 2015). The equation to calculate change in pressure head across the system is:

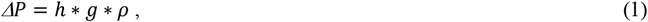

where *ΔP* is the pressure (Pa) of the system; *h* is the height of the water reservoir (m); *g* is the acceleration due to gravity 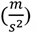, and *ρ* is the temperature dependent density of the water 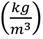

The equation for mass flowrate is:

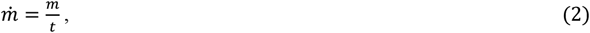

where 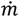 is the mass flowrate 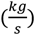; *m* is the weight of the water on the balance (kg), and *t* is the time (s).

The hydraulic conductance calculation combines the mass flow rate, and the pressure as follows:

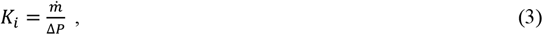

Where *K*_*i*_ if the ith hydraulic conductance (10^−6^ s * m); 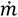 is the mass flowrate on the precision balance 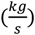 from Equation 2, and Δ*P* is the pressure head across the system from Equation 1 in (MPa).

The specific hydraulic conductivity (Ks) was also computed. The difference between specific hydraulic conductivity and hydraulic conductivity is that the stem length and xylem cross-sectional area of the samples are used to normalize the conductance. The unit of the hydraulic conductivity is expressed differently across articles (Pérez-Harguindeguy et al., 2013), therefore, we chose the irreducible unit representation. The equation of Ks is:

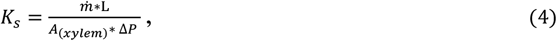

Where *K*_*s*_ is the Initial Specific Hydraulic Conductivity, (10^−6^ * *s*); 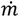 is the mass flowrate on the precision balance 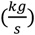 from Equation 2; *L* is the length of the sample, and Δ*P* is the pressure head difference across the system from Equation 1 in MPa. The typical unit representation for *K*_*s*_ in the literature is (kg *m*^−1^ *s*^−1^ *MPa*^−1^) (Choat et al., 2015; Pérez-Harguindeguy et al., 2013; J. S. Sperry & Tyree, 1990). (kg *m*^−1^ *s*^−1^ *MPa*^−1^) reduces to the simpler form of (10^−6^ * *s)* which we use here.

Any naturally occurring embolisms are removed and the maximum specific conductivity is measured after the natural specific conductivity is measured. The stem sample was placed into a tank partially filled with DI water and the headspace of the tank was de-pressurized to -90kPa. Samples were kept in the tank for overnight. This method was shown to be effective by (Sperry et al., 1988). The procedures and calculations outlined above are repeated on this flushed stem section to obtain the theoretical max specific hydraulic conductivity. This maximum conductivity value is the basis of comparison for the percentage loss of conductance (PLC) of a stem.

Embolies were reintroduced at incrementally increasing pressure values to simulate water stress in the stem sample. Synthetic water stress was induced by the cavitation chamber. The ‘air seeding’ method was used to induce embolism and Sperry & Tyree by pushing air into a xylem using a Scholander pressure chamber (Sperry & Tyree, 1990). The cavitation chamber applies nitrogen gas into the stems. In this way, the xylem was partially filled by nitrogen and a synthetic water stress occurred with embolism in xylem (Sheridan & Nackley, 2021).

The measurement protocol above and associated calculations were again repeated through a set of increasing pressures until the conductivity of the stem is zero. We took higher resolution (in pressure increments) than typical measurements to increase the confidence in the observed differences in the shape of the resulting vulnerability curve. Note that in this manuscript ‘vulnerability curve’ is used to describe the functional shape and fitted function to the data, whereas ‘PLC’ is used to describe the axes numerical value of the loss in conductance. Typical PLC graphics are drawn with five pressure applications. However, we applied pressures to the samples up to ten times resulting in a more highly resolved vulnerability curve. All stems (sixteen) were exposed to the same pressures. These given pressures were 0.15Mpa, 0.25MPa, 0.50Mpa, 0.75MPa, 1.0MPa, 1.25MPa, 1.50MPa, 1.75MPa, 2.0MPa, 2.25Mpa. All stems have been exposed to each pressure for 10 minutes. After the cavitation chamber, stems have been put in the tray which is filled with DI water. The aim of this step is to allow stems find the equilibrium (Sperry & Tyree, 1990). When this step was omitted, it was observed that the stem sprayed the air bubbles from xylem and filled the connection parts of the apparatus with air bubbles. A photograph of the cavitation chamber and associated components is shown in Figure 6.

**Figure 6.**
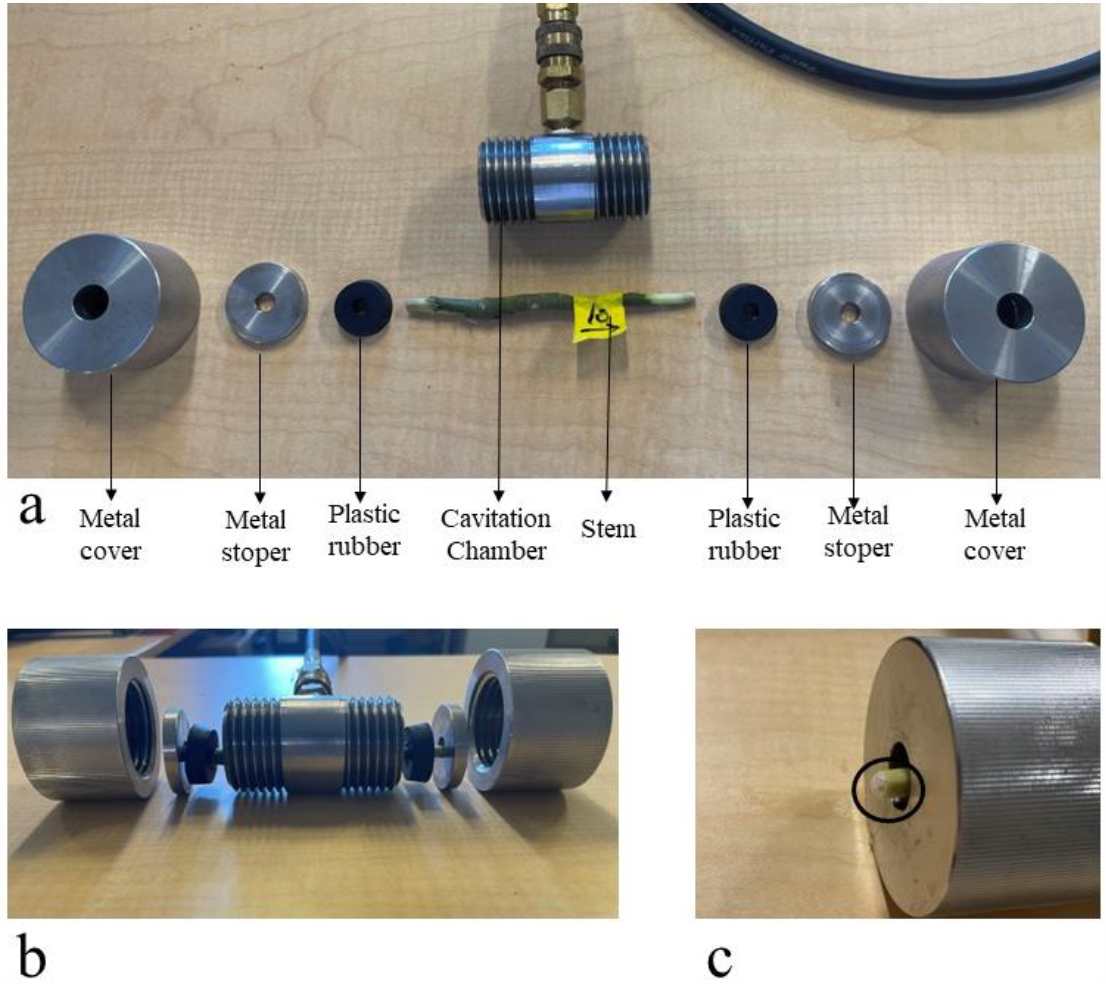
a) all parts of the cavitation chamber, b) assembling parts of the cavitation chamber and c) bubbles coming out of the xylem thanks to applied pressure

The aim of the study is to understand percentage loss of hydraulic conductivity in plant stems. The formula of the percentage loss of conductance is:

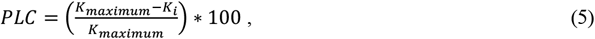

Where *PLC* is the percentage loss of conductance (%); *K*_*maximum*_ is the maximum hydraulic conductivity where after all embolies were removed (10^−6^ * *s*); *K*_*i*_ is the specific hydraulic conductivity after pressure, i, was applied (10^−6^ * *s*). Here, i, is a value in the set of 0.15Mpa, 0.25MPa, 0.50Mpa, 0.75MPa, 1.0MPa, 1.25MPa, 1.50MPa, 1.75MPa, 2.0MPa, and 2.25Mpa.

A two-tailed t-test, regression analysis and dimensional analysis were used in this study. A two-tailed t-test was used to understand which measurable value is important for our analysis. All quantifiable parts of the samples, including length, xylem area, diameter, and pith area, were taken into consideration and the differences in both the mean of these values and p-values were observed in the two separated groups. A detailed explanation of the results is shown in the results section. All two-tailed t-test analysis was done in SPSS software. Our research’s regression analysis forced the intercept through 0. This is because if there is not any pressure which applies to our samples, there cannot be any hydraulic conductivity losses. Regression analyses were performed in Python.

## 2. Result

All data were imported into tables.Table 1 and Table 2 show all collected and calculated data before the cavitation chamber measurements.

**Table 1.**
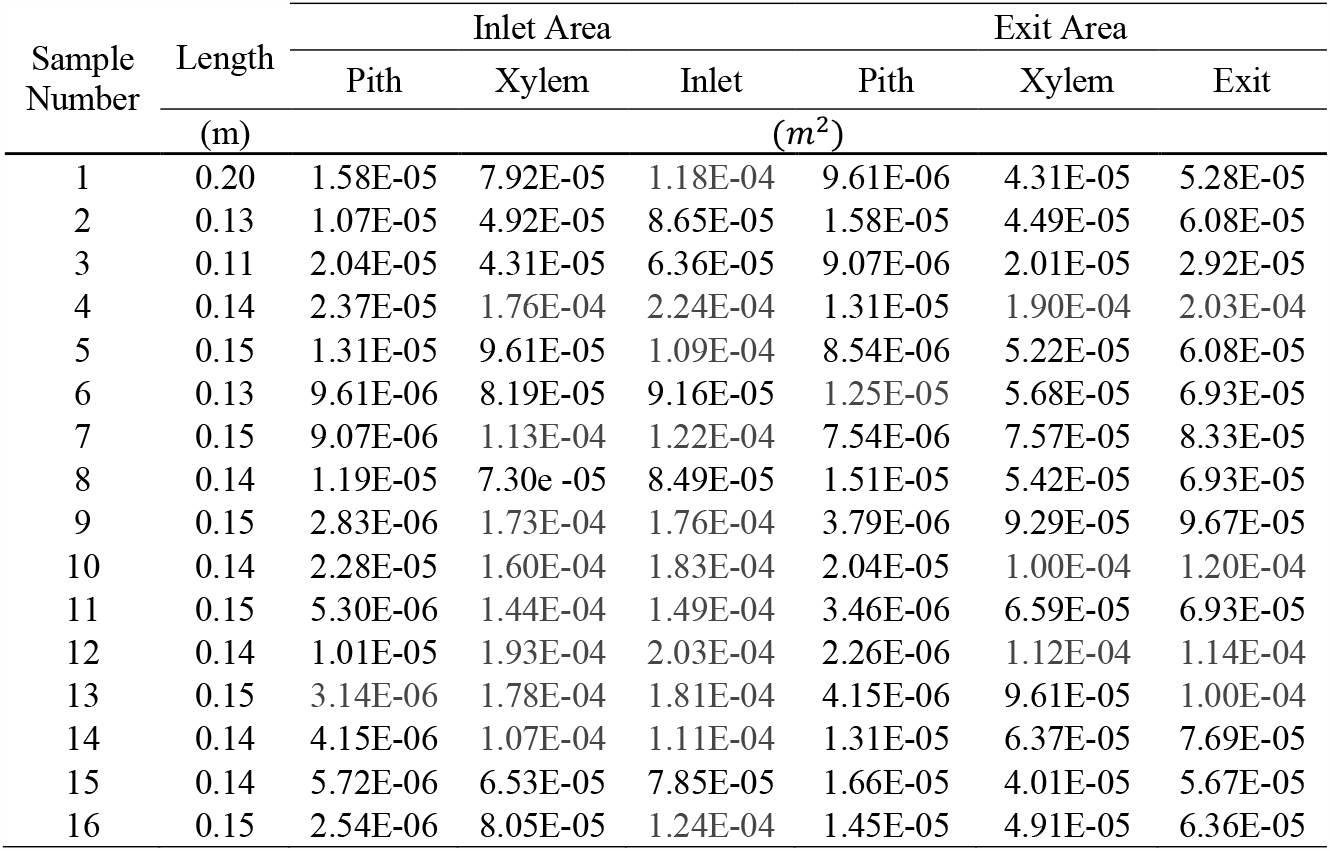
Geometric features of the samples.

| Sample Number | Length<br>(m) | Inlet Area |  |  | Exit Area |  |  |
| --- | --- | --- | --- | --- | --- | --- | --- |
|  |  | Pith | Xylem | Inlet | Pith | Xylem | Exit |
|  |  | (m<sup>2</sup>) |  |  |  |  |  |
| 1 | 0.20 | 1.58E-05 | 7.92E-05 | 1.18E-04 | 9.61E-06 | 4.31E-05 | 5.28E-05 |
| 2 | 0.13 | 1.07E-05 | 4.92E-05 | 8.65E-05 | 1.58E-05 | 4.49E-05 | 6.08E-05 |
| 3 | 0.11 | 2.04E-05 | 4.31E-05 | 6.36E-05 | 9.07E-06 | 2.01E-05 | 2.92E-05 |
| 4 | 0.14 | 2.37E-05 | 1.76E-04 | 2.24E-04 | 1.31E-05 | 1.90E-04 | 2.03E-04 |
| 5 | 0.15 | 1.31E-05 | 9.61E-05 | 1.09E-04 | 8.54E-06 | 5.22E-05 | 6.08E-05 |
| 6 | 0.13 | 9.61E-06 | 8.19E-05 | 9.16E-05 | 1.25E-05 | 5.68E-05 | 6.93E-05 |
| 7 | 0.15 | 9.07E-06 | 1.13E-04 | 1.22E-04 | 7.54E-06 | 7.57E-05 | 8.33E-05 |
| 8 | 0.14 | 1.19E-05 | 7.30E-05 | 8.49E-05 | 1.51E-05 | 5.42E-05 | 6.93E-05 |
| 9 | 0.15 | 2.83E-06 | 1.73E-04 | 1.76E-04 | 3.79E-06 | 9.29E-05 | 9.67E-05 |
| 10 | 0.14 | 2.28E-05 | 1.60E-04 | 1.83E-04 | 2.04E-05 | 1.00E-04 | 1.20E-04 |
| 11 | 0.15 | 5.30E-06 | 1.44E-04 | 1.49E-04 | 3.46E-06 | 6.59E-05 | 6.93E-05 |
| 12 | 0.14 | 1.01E-05 | 1.93E-04 | 2.03E-04 | 2.26E-06 | 1.12E-04 | 1.14E-04 |
| 13 | 0.15 | 3.14E-06 | 1.78E-04 | 1.81E-04 | 4.15E-06 | 9.61E-05 | 1.00E-04 |
| 14 | 0.14 | 4.15E-06 | 1.07E-04 | 1.11E-04 | 1.31E-05 | 6.37E-05 | 7.69E-05 |
| 15 | 0.14 | 5.72E-06 | 6.53E-05 | 7.85E-05 | 1.66E-05 | 4.01E-05 | 5.67E-05 |
| 16 | 0.15 | 2.54E-06 | 8.05E-05 | 1.24E-04 | 1.45E-05 | 4.91E-05 | 6.36E-05 |

**Table 2.** All calculated data after hydraulic conductivity measurement apparatus.

| Sample Number | Pressure Head | Flow Rate | Native Specific Conductivity | Maximum Specific Conductivity | Stem Water Potential |
| --- | --- | --- | --- | --- | --- |
|  | (m) | (kg/s) | 10<sup>-6</sup> * s | 10<sup>-6</sup> * s | (kPa) |
| 1 | 0.83 | 3.33E-07 | 0.10 | 0.18 | -200 |
| 2 | 0.83 | 3.66E-07 | 0.12 | 0.46 | -250 |
| 3 | 0.82 | 5.33E-07 | 0.18 | 0.23 | -500 |
| 4 | 0.82 | 1.00E-07 | 0.01 | 0.08 | -500 |
| 5 | 0.83 | 5.66E-07 | 0.11 | 0.21 | -700 |
| 6 | 0.82 | 4.33E-07 | 0.08 | 0.39 | -700 |
| 7 | 0.82 | 2.20E-06 | 0.36 | 0.43 | -850 |
| 8 | 0.82 | 2.46E-06 | 0.62 | 0.71 | -750 |
| 9 | 0.82 | 9.66E-07 | 0.10 | 0.19 | -550 |
| 10 | 0.82 | 3.00E-06 | 0.32 | 0.43 | -600 |
| 11 | 0.82 | 4.33E-07 | 0.05 | 0.20 | -500 |
| 12 | 0.82 | 8.66E-07 | 0.07 | 0.14 | -500 |
| 13 | 0.82 | 4.33E-07 | 0.04 | 0.10 | -350 |
| 14 | 0.82 | 2.00E-07 | 0.03 | 0.10 | -350 |
| 15 | 0.82 | 3.33E-07 | 0.10 | 0.17 | -400 |
| 16 | 0.82 | 3.66E-07 | 0.09 | 0.16 | -400 |

The average difference in specific conductivity (natural vs. max) across all samples was found to be 42.67%. We inferred that even though all plants were irrigated in same schedule to minimize water stress, emboli levels were non-zero and variable across the sample set. A two-tailed t-test of the mean difference resulted in a p-value of 0.004 indicating that the conductivity was increased by a statistically significant amount after emboli removal.

The PLC values for all 16 samples are presented in Figure. Here it is evident that each stem can have distinct responses to stress. Therefore, underlying variables that may contribute to these alternative responses are investigated below.

### 2.1 Statistical Analysis Results

#### 2.1.1 Xylem average area

Two-tailed t-tests were performed in SPSS software program to understand the significance of average xylem area. All samples were classified to two different group according to their vulnerability curve shape (seen in Figure 8). Average xylem was calculated because samples had different xylem areas for both inlet and exit sections. Therefore, taking the average of xylem area can be more accurate for our results. The two-tailed t-test of the mean difference resulted in a p-value of 0.003 indicating that the average xylem area is quite significant for our results.

A Mann-Whitney U test was applied to the same grouping of our data. The Mann Whitney U test of the mean difference resulted in a p-value of 0.019 indicating that the average xylem area is quite significant according to both two-tailed t-test and Mann Whitney U test.

#### 2.1.2 Length of the samples

The statistical analysis described above was repeated for stem length. Here, the two-tailed t-test of the mean difference resulted in a p-value of 0.990, the Mann-Whitney U test resulted in a p-value of 0.441. These p-values indicate that the length of the samples is not significant for our results.

### 2.2 The relation between xylem area and vulnerability curve

The Figure8 again presents the PLC values of all samples, but now each point is colored by the individual stem’s average xylem cross sectional area. Here there appears to be 2 groups, thinner stems where the vulnerability curve is not linear and concave in shape and stems with more xylem having a near linear vulnerability response. Stated another way, the samples which have smallest area lost 90% of their hydraulic conductivity after at ∼1 MPa; however, the samples with the largest area have lost only ∼50% of their hydraulic conductivity at the same pressure levels.

### 2.3 The relation between sample length and vulnerability curve

Figure9 presents the PLC values of all samples, but now each point is colored by the individual stem’s length. Here, any potential relationship is less apparent. This is consistent with a prior study that shows that short and long segments gave similar conductivity results. (Choat et al., 2015; Cochard et al., 2010).

### 2.4 Log-log Graphic between PLC and Pressures for all samples

The log-log graph is used to investigate if the relationship between the x and y axes follows a power law relationship. A power law relationship is likely if the lines in the log-log plot are linear (straight lines). Straight lines were obtained as anticipated when the results of the log-log graphic were examined, in Figure10. As a result, we will seek to fit the data with a power law function in the next sections. The data in Figure 10 also suggest that the exponent for each stem could be a function of the stem’s physical properties. That is, each stem has a unique slope in the log-log plot, therefore each stem would have a unique exponent in a power law relationship.

### 2.5 PLC graphic with xylem pressures

Our goal was to seek a more universal and unified equation for the vulnerability curve. The log-log plot (Figure10) suggests a power law functional for, but the exponent of the power law appears to be related to the xylem cross sectional area. Therefore, we adopt our fit such that the exponent in the power law is a function of stem properties. More specifically, and due to earlier findings, we select the average xylem area (pressure) as an additional independent variable. Our choice of xylem area is supported by the statistical analysis presented above.

We also seek a dimensionless expression within these constraints and arrive at the following functional fit (vulnerability curve) to the PLC values.

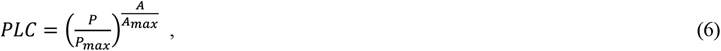

where *PLC* is the Percentage of Loss Hydraulic Conductivity (Observed Values) in percent; *P* is the pressure applied to the samples (MPa); *P*_*max*_ is the pressure where the conductivity of the stems is thought to be completely lost (constant =2.4MPa); *A* is the average cross-sectional area of the stem, and *A*_*max*_ is the maximum cross-sectional area (constant =0.000183473 m^2^). Note that equation 6 is only defined for Pressures P<P_max_ and average cross sectional areas A<A_max_. The regression of the function presented in Equation 6 against all measurements is shown I Figure 11, and the *r*^*2*^ value for this fit was calculated as 0.74.

## 3. Discussion

Direct comparisons between plants across the literature can be challenging because of the diversity in plants, plant age, and cavitation approach. There are, however, studies that compare the sensitivity of different plants directly (summarized in Table 3).

Table 3 provides a comparison of P_min_ and Pmax values from previous studies on eight different plant species. Among these plants, hemp and Malus domestica borkh exhibit the most vulnerable stems under water stress, indicated by their lowest Pmax values. Hemp also shows an early response threshold to stress, as reflected by its minimal P_min_ value. The shape of the vulnerability curve is another crucial aspect of vulnerability curve research. Previous studies have reported a concave curve for Abies balsama and Picea Rubens, indicating an increased steepness with higher pressure. In contrast, a sigmoidal curve, the expected shape, was observed for Sugar maple, Juniperus virginiana, and Malus var. While Malus domestica displayed a sigmoidal curve, it was nearly linear. Laurus nobilis exhibited a convex curve, becoming less steep with higher pressure. Notably, both hemp and Laurus nobilis had low Pmax values and convex vulnerability curves. Furthermore, hemp and Malus domestica borkh showed potential expressions of vulnerability curves that were close to linear, resembling a power law. These non-sigmoidal vulnerability curve expressions seem to be associated with plants having lower Pmax values. Conversely, the four plants with the highest Pmax values all expressed sigmoidal vulnerability curves, suggesting a potential link between Pmax value and vulnerability curve shape.

Within our hemp study, we observed a variation in the shape of the vulnerability curve, ranging from highly convex to near-linear (Figure 7). This led us to hypothesize that an additional independent variable, aside from pressure, contributes to or partially explains this shape change. To explore this further, we categorized the vulnerability curves into two groups based on their shape: highly convex and near-linear. We then examined other independent variables associated with each curve, such as xylem area and stem length. By performing a two-tailed t-test, we found that the average xylem area showed significant differences between the groups at a confidence level of 95% (p-value = 0.003). However, the t-test for stem length resulted in a p-value of 0.99, indicating that the length of the samples was not significant. These stem length findings align with previous studies (Choat et al., 2015; Cochard et al., 2010).

**Figure 7.**
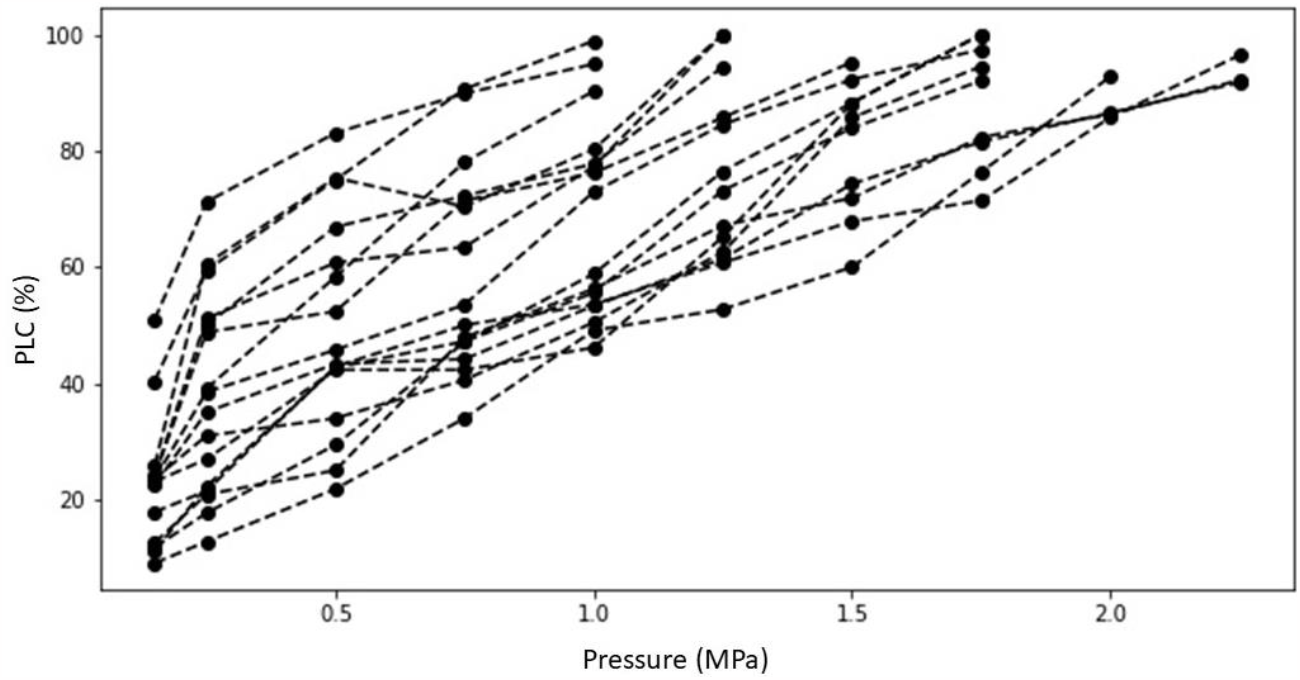
PLC graphics for all samples.

Figure 8 visually represents these findings, with curves colored according to the average xylem area and associated with the vulnerability curve shape groups. In contrast, Figure 9 shows the curves colored by stem length, where the coloring does not align with the vulnerability curve shape groups. It can be observed in Figure 8 that stems with smaller xylem areas exhibit more nonlinear (convex) vulnerability curve behavior, while stems with larger xylem areas have near-linear curves. Convex, concave, and linear curves can all be mathematically described by a power law, with the exponent determining the shape. Assuming a power law behavior, thin stems would have an exponent less than 1 (convex), while thicker stems would have an exponent near 1 (linear). Note that an exponent greater than 1 would result in a concave shape. The simplest and most parsimonious mathematical expression to describe this behavior would relate the exponent to a normalized area, limited to stem areas less than the maximum area defined in Equation 6.

**Figure 8.**
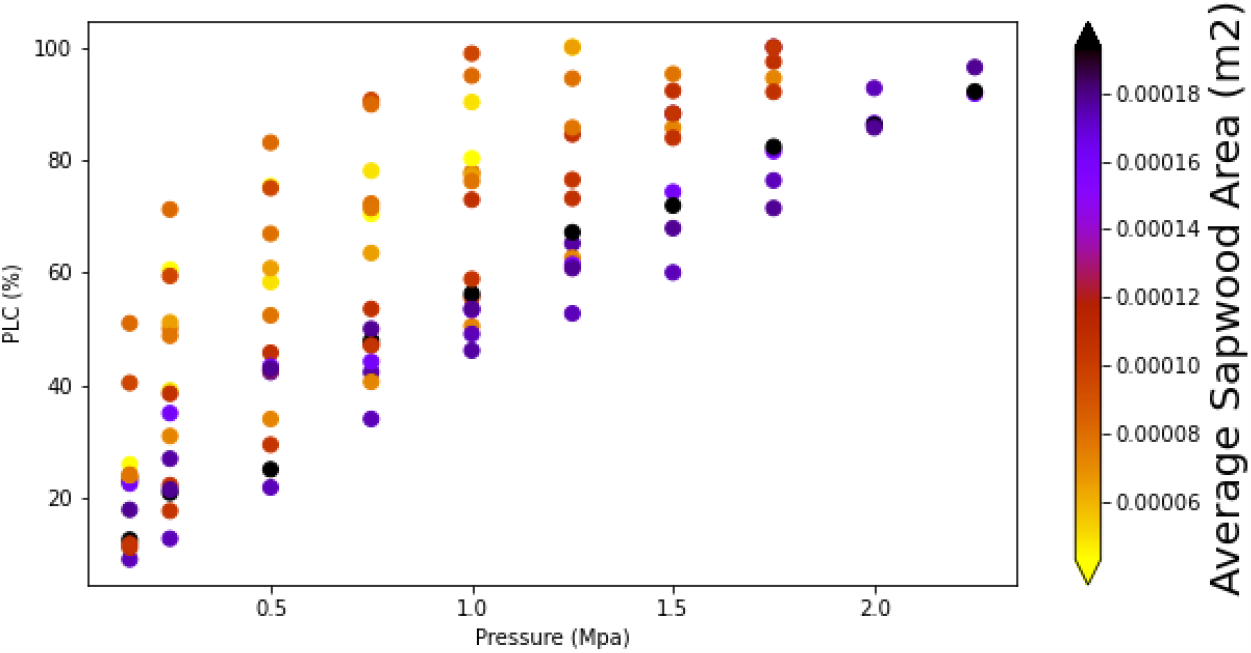
Illustrates the vulnerability curves with average sapwood areas.

**Figure 9.**
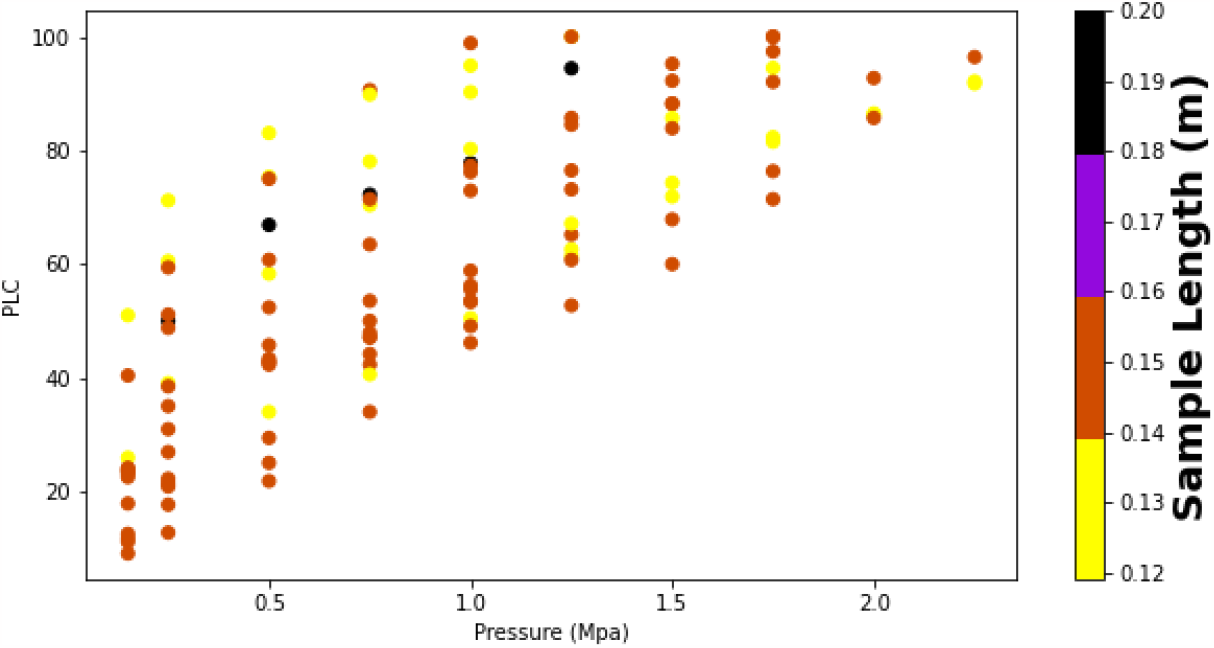
Illustrates the vulnerability curves with colored by sample length.

**Figure 10.**
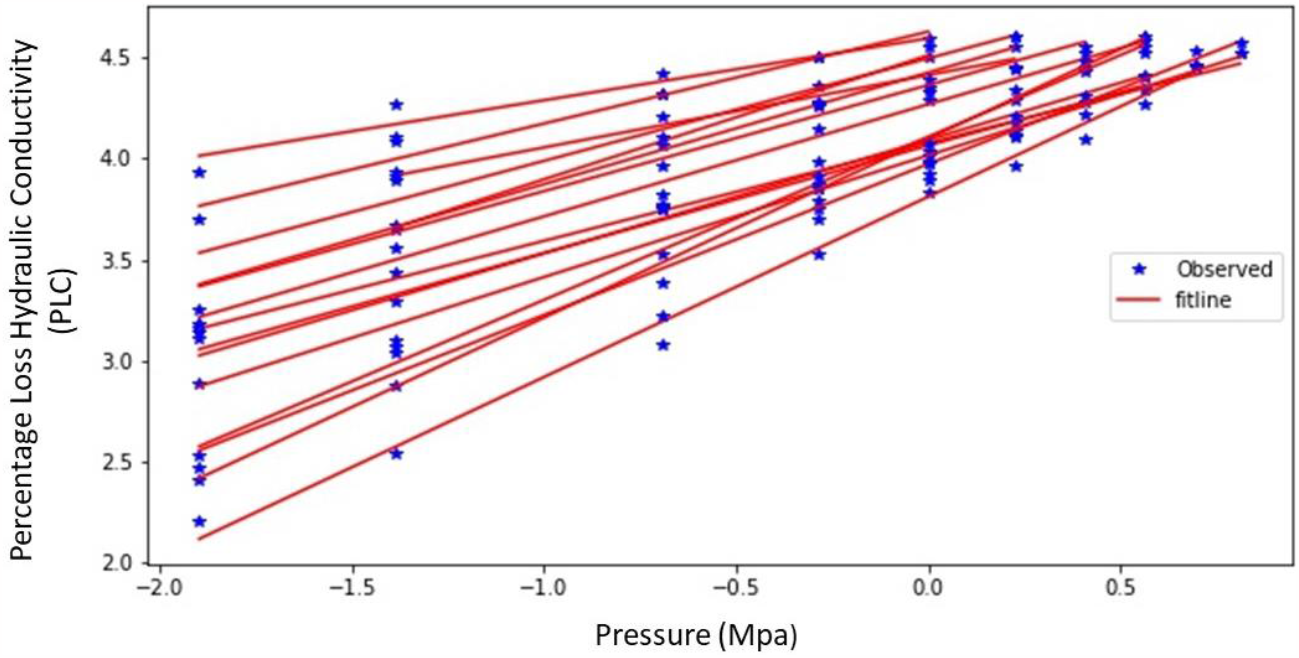
Log-log graphic between pressures and PLC

**Figure 11.**
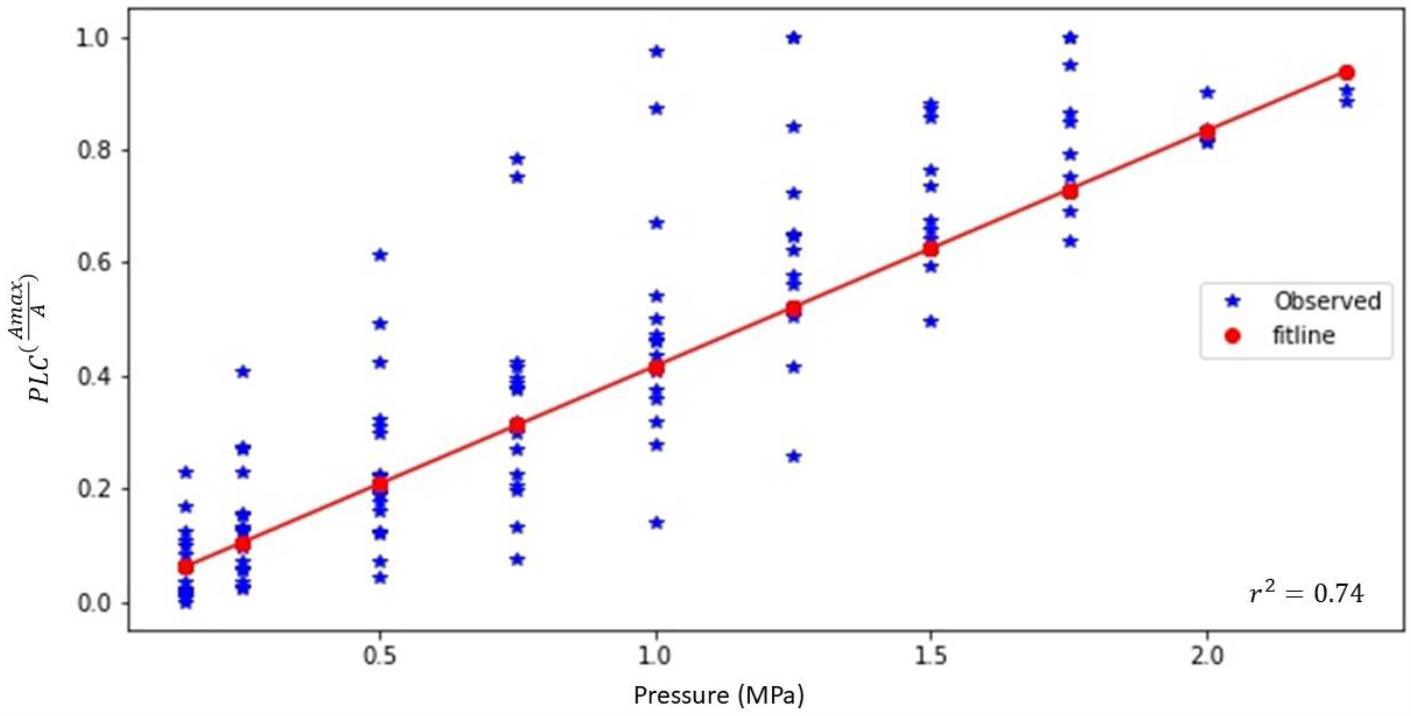
Percentage loss of hydraulic conductivity of hemp stems under water stress conditions.

Finally, a power law function may also be applicable to describe concave vulnerability curves. Although we did not observe concave vulnerability curves in our hemp data, they have been reported in other studies (see Table 3). We hypothesize that a power law may be sufficient to describe vulnerability curve shapes that are not sigmoidal. However, further in-depth analysis and meta-analysis would be necessary to test this hypothesis against observations. Nevertheless, the power law proved effective in capturing the highly variable shapes observed in our hemp study.

**Table 3.** Comparing the maximum and minimum pressure differences around some plants.

| Plant | P<sub>min</sub><br>(MPa) | P<sub>max</sub><br>(MPa) | Vulnerability<br>Shape | Reference |
| --- | --- | --- | --- | --- |
| Hemp | 0.15 | 2.4 | Power Law | This study |
| Malus domestica<br>borkh | 0.25 | 2.4 | Sigmoidal/linear | (De Baerdemaeker et<br>al., 2019) |
| Sugar maple | 0.30 | 4.5 | Sigmoidal | (Sperry & Tyree,<br>1988) |
| Laurus nobilis | 0.5 | 3.0 | Convex | (Hacke & Sperry,<br>2003) |
| Malus var. | 0.5 | 6.0 | Sigmoidal | (Sheridan & Nackley,<br>2021) |
| Sugar maple | 0.5 | 5.0 | Sigmoidal | (Melcher, Zwieniecki,<br>& Holbrook, 2003) |
| Picea rubens | 1 | 4.2 | Concave | (Sperry & Tyree,<br>1990) |
| Abies balsamea | 1 | 3.5 | Concave | (Sperry & Tyree,<br>1990) |
| Juniperus<br>virginiana | 2 | 8.0 | Sigmoidal | (Sperry & Tyree,<br>1990) |

## 4. Conclusion

The aim of this study was to investigate the vulnerability curve of hemp stems under water stress conditions, providing valuable insights into the plant’s response to water scarcity and informing water management practices. Hemp was chosen as the focus of this research due to its rapid expansion as an agricultural commodity and the limited knowledge about its vulnerability curve. The methodology involved inducing embolism in the xylem using the ‘air seeding’ method and measuring the stem hydraulic conductivity at various stress states using a gravity hydraulic flow apparatus. These measurements were then used to construct the vulnerability curve for hemp, capturing its response to drought conditions.

The vulnerability curve of hemp was compared with those of other plants studied in previous research. The findings indicated that hemp exhibited greater sensitivity to water stress compared to other plants. The onset of stress occurred at lower pressures, and even small pressure changes led to significant loss of hydraulic conductivity in hemp xylem. Additionally, the pressure at which stems lost 90% of their conductivity was the lowest among the plants studied, suggesting a higher vulnerability to water stress in hemp. This highlights the importance of adequate watering for hemp cultivation to mitigate water stress effects. The shape of the vulnerability curve for hemp was influenced by stem geometry and size. Thinner and younger stems with smaller xylem area demonstrated higher sensitivity to water stress, as depicted in the PLC graphic (Figure 8). Statistical analysis using a two-tailed t-test supported this observation, showing a significant difference in average xylem area between groups (p-value=0.003 at 95% confidence interval). Consequently, a mathematical equation incorporating both average xylem area and pressure was formulated to fit the data, with a simple power law (Equation 6) explaining over 70% of the variance in the measurements.

While conventional literature suggests that vulnerability curves tend to exhibit a sigmoidal shape with respect to pressure, this study found that hemp displayed a non-sigmoidal shape. Instead, a multivariate power law model proved to be more effective and parsimonious in capturing the vulnerability curve for hemp. It should be noted that non-sigmoidal vulnerability curves, including convex and concave shapes, have been reported for other plant species. Further research and meta-analysis are required to explore the potential of these variations in water stress response across different plant phenotypes.

This study presents an opportunity for further investigation specific to hemp. Building upon the findings of this research, future studies can focus on growing hemp under regulated deficit conditions and comparing the vulnerability curves of plants experiencing water stress with those of well-irrigated plants. This approach would provide insights into the differences in response to maximum drought stress between the two groups and allow for a comprehensive understanding of hemp’s water stress tolerance. In summary, this study contributes to our understanding of the vulnerability curve for hemp, highlighting its higher sensitivity to water stress compared to other plants. The findings underscore the importance of proper watering practices in hemp cultivation. Stem geometry and size were identified as key factors shaping the vulnerability curve, with thinner and younger stems exhibiting greater sensitivity to water stress. The use of a multivariate power law model proved effective in describing the vulnerability curve for hemp. Further research is needed to explore non-sigmoidal vulnerability curves and expand studies specific to hemp under various water stress conditions.

## Author Contributions

Conceptualization, C.H., H.A.A., L.N. and M.Y; methodology, M.Y; software, M.Y; validation, M.Y; formal analysis, C.H., H.A.A. and M.Y; investigation, M.Y; resources, C.H., H.A.A. and L.N.; data curation, M.Y; writing—original draft preparation, M.Y., H.A.A. and C.H.; writing—review and editing, M.Y, H.A.A., C.H. and L.N.; visualization, C.H., H.A.A. and M.Y; supervision, C.H., H.A.A. and L.N.; project administration, C.H.; funding acquisition, C.H. All authors have read and agreed to the published version of the manuscript.

## Data Availability Statement

No Data attached.

## Acknowledgments

I would like to think of the General Directorate of Agricultural Research and Policies, and Republic of Türkiye Ministry of Agriculture and Forestry for supporting this project.

## References

Afrin, F., Chi, M., Eamens, A. L., Duchatel, R. J., Douglas, A. M., Schneider, J., … Dun, M. D. (2020). cancers Can Hemp Help? Low-THC Cannabis and Non-THC Cannabinoids for the Treatment of Cancer. https://doi.org/10.3390/cancers12041033

Al-agele, H. A., Jashami, H., & Higgins, C. W. (2022). Evaluation of novel ultrasonic sensor actuated nozzle in center pivot irrigation systems. Agricultural Water Management, 262, 107436.

AL-agele, H. A., Jashami, H., Nackley, L., & Higgins, C. (2021). A variable rate drip irrigation prototype for precision irrigation. Agronomy, 11(12), 2493.

Al-agele, H. A., Mahapatra, D. M., Prestwich, C., & Higgins, C. W. (2020). Dynamic Adjustment of Center Pivot Nozzle Height: An Evaluation of Center Pivot Water Application Pattern and the Coefficient of Uniformity. Applied Engineering in Agriculture, 36(5), 647–656.

Al-Agele, H. A., Nackley, L., & Higgins, C. (2021a). Testing novel new drip emitter with variable diameters for a variable rate drip irrigation. Agriculture (Switzerland), 11(2), 1–8. https://doi.org/10.3390/agriculture11020087

Al-Agele, H. A., Nackley, L., & Higgins, C. W. (2021b). A pathway for sustainable agriculture. Sustainability (Switzerland), 13(8). https://doi.org/10.3390/su13084328

Anjum, S. A., Xie, X. yu, Wang, L. chang, Saleem, M. F., Man, C., & Lei, W. (2011). Morphological, physiological and biochemical responses of plants to drought stress. African Journal of Agricultural Research, 6(9), 2026–2032. https://doi.org/10.21921/jas.5.3.7

Brodribb, T. J. (2009). Xylem hydraulic physiology: The functional backbone of terrestrial plant productivity. Plant Science, 177(4), 245–251. https://doi.org/10.1016/j.plantsci.2009.06.001

Bucci, S. J., Scholz, F. G., Goldstein, G., Meinzer, F. C., Da, & L., & Sternberg, S. L. (2003). Dynamic changes in hydraulic conductivity in petioles of two savanna tree species: factors and mechanisms contributing to the refilling of embolized vessels. Plant, Cell and Environment (Vol. 26).

Choat, B., Creek, A. D., Lo Gullo, M. A., Nardini, A., Oddo, E., Raimondo, F., … Vilagrosa, A. (2015). PrometheusWiki - Quantification of vulnerability to xylem embolism - Bench dehydration Quantification of vulnerability to xylem embolism - bench dehydration, (MAY).

Choné, X., Van Leeuwen, C., Dubourdieu, D., & Gaudillère, J. P. (2001). Stem water potential is a sensitive indicator of grapevine water status. Annals of Botany, 87(4), 477–483. https://doi.org/10.1006/anbo.2000.1361

Cochard, H., Badel, E., Herbette, S., Delzon, S., Choat, B., & Jansen, S. (2013). Methods for measuring plant vulnerability to cavitation: A critical review. Journal of Experimental Botany. https://doi.org/10.1093/jxb/ert193

Cochard, H., Herbette, S., Barigah, T., Badel, E., Ennajeh, M., & Vilagrosa, A. (2010). Does sample length influence the shape of xylem embolism vulnerability curves? A test with the Cavitron spinning technique. Plant, Cell and Environment, 33(9), 1543–1552. https://doi.org/10.1111/j.1365-3040.2010.02163.x

De Baerdemaeker, N. J. F., Arachchige, K. N. R., Zinkernagel, J., Van Den Bulcke, J., Van Acker, J., Schenk, H. J., … Tognetti, R. (2019). The stability enigma of hydraulic vulnerability curves: Addressing the link between hydraulic conductivity and drought-induced embolism. Tree Physiology, 39(10), 1646–1664. https://doi.org/10.1093/treephys/tpz078

De Silva, N. D. G., Cholewa, E., & Ryser, P. (2012). Effects of combined drought and heavy metal stresses on xylem structure and hydraulic conductivity in red maple (Acer rubrum L.). Journal of Experimental Botany, 63(16), 5957–5966. https://doi.org/10.1093/jxb/ers241

Fereres, E., & Soriano, M. A. (2007). Deficit irrigation for reducing agricultural water use. In Journal of Experimental Botany (Vol. 58, pp. 147–159). https://doi.org/10.1093/jxb/erl165

García-Tejero, I. F., Durán-Zuazo, V. H., Pérez-Álvarez, R., Hernández, A., Casano, S., Morón, M., & Muriel-Fernández, J. L. (2014). Impact of plant density and irrigation on yield of hemp (Cannabis sativa L.) in a mediterranean semi-arid environment. Journal of Agricultural Science and Technology, 16(4), 887–895.

Gill, A. R., Loveys, B. R., Cowley, J. M., Hall, T., Cavagnaro, T. R., & Burton, R. A. (2022). Physiological and morphological responses of industrial hemp (Cannabis sativa L.) to water deficit. Industrial Crops and Products, 187(PA), 115331. https://doi.org/10.1016/j.indcrop.2022.115331

Gleason, S. M., Wiggans, D. R., Bliss, C. A., Comas, L. H., Cooper, M., DeJonge, K. C., … Zhang, H. (2017). Coordinated decline in photosynthesis and hydraulic conductance during drought stress in Zea mays. Flora: Morphology, Distribution, Functional Ecology of Plants, 227, 1–9. https://doi.org/10.1016/J.FLORA.2016.11.017

Hacke, U. G., & Sperry, J. S. (2003). Limits to xylem refilling under negative pressure in Laurus nobilis and Acer negundo. Plant, Cell and Environment, 26(2), 303–311. https://doi.org/10.1046/j.1365-3040.2003.00962.x

Hargrave, K. R., Kolb, K. J., Ewers, F. W., & Davis, S. D. (1994). Conduit diameter and drought-induced embolism in Salvia mellifera Greene (Labiatae). New Phytologist, 126(4), 695–705. https://doi.org/10.1111/j.1469-8137.1994.tb02964.x

Holbrook, N. M., & Zwieniecki, M. A. (2005). Vascular Transport in Plants.

Holtzman, N. M., Anderegg, L. D. L., Kraatz, S., Mavrovic, A., Sonnentag, O., Pappas, C., … Konings, A. G. (2021). L-band vegetation optical depth as an indicator of plant water potential in a temperate deciduous forest stand. Biogeosciences, 18(2), 739–753. https://doi.org/10.5194/bg-18-739-2021

Irmak, S., Odhiambo, L. O., Kranz, W. L., & Eisenhauer, D. E. (2011). Irrigation efficiency and uniformity, and crop water use efficiency.

Jones, H. G. (1990). Physiological Aspects of the Control of Water Status in Horticultural Crops. HortScience, 25(1), 19–25. https://doi.org/10.21273/hortsci.25.1.19

Khokhar, T. (2017, March). Chart: Globally, 70% of Freshwater is Used for Agriculture.

Li, H.-L. (1974). An Archaeological and Historical Account of Cannabis in China (Vol. 28).

Lovisolo, C., & Schubert, A. (1998). Effects of water stress on vessel size and xylem hydraulic conductivity in Vitis vinifera L. Journal of Experimental Botany, 49(321), 693–700. https://doi.org/10.1093/JXB/49.321.693

McCutchan, H., & Shackel, K.. (1992). Stem-water Potential as a Sensitive Indicator of Water Stress in Prune Trees (Prunus domestica L. cv. French). Journal of the American Society for Horticultural Science, 117(4), 5.

Melcher, P. J., Zwieniecki, M. A., & Holbrook, N. M. (2003). Vulnerability of xylem vessels to cavitation in sugar maple. Scaling from individual vessels to whole branches. Plant Physiology, 131(4), 1775–1780. https://doi.org/10.1104/pp.102.012856

Oki, T., & Kanae, S. (2006). Global hydrological cycles and world water resources. Science, 313(5790), 1068–1072. https://doi.org/10.1126/science.1128845

Onder, S., Emin, M., & Onder, D. (2005). Different irrigation methods and water stress effects on potato yield and yield components, 73, 73–86. https://doi.org/10.1016/j.agwat.2004.09.023

Osakabe, Y., Osakabe, K., Shinozaki, K., & Tran, L. S. P. (2014). Response of plants to water stress. Frontiers in Plant Science, 5(MAR). https://doi.org/10.3389/fpls.2014.00086

Pérez-Harguindeguy, N., Díaz, S., Garnier, E., Lavorel, S., Poorter, H., Jaureguiberry, P., … Cornelissen, J. H. C. (2013). New handbook for standardised measurement of plant functional traits worldwide. Australian Journal of Botany, 61(3), 167–234. https://doi.org/10.1071/BT12225

Qin, D., Qian, Y., Han, L., Wang, Z., Li, C., & Zhao, Z. (2011). Assessing impact of irrigation water on groundwater recharge and quality in arid environment using CFCs, tritium and stable isotopes, in the Zhangye Basin, Northwest China. Journal of Hydrology, 405(1–2), 194–208. https://doi.org/10.1016/j.jhydrol.2011.05.023

Schlosser, C. A., Strzepek, K., Gao, X., Gueneau, A., Fant, C., Paltsev, S., … Reilly, J. (2014). The Future of Global Water Stress: An Integrated Assessment.

Scholander, P. F., Hammel, H. T., Bradstreet, E. D., & Hemmingsen, E. A. (1965). Sap pressure in vascular plants. Science (Vol. 148). https://doi.org/10.1126/science.148.3668.339

Shao, H. B., Chu, L. Y., Jaleel, C. A., & Zhao, C. X. (2008). Water-deficit stress-induced anatomical changes in higher plants. Comptes Rendus - Biologies, 331(3), 215–225. https://doi.org/10.1016/j.crvi.2008.01.002

Sheridan, R. A., & Nackley, L. L. (2021). A Primer on Plant Hydraulic Physiology for Nursery Professionals. Tree Planters’ Notes, 64(2), 70–79.

Smathers, R. L., King, B. A., & Patterson, P. E. (1995). Economics of Surface Irrigation Systems, (779), 16.

Sperry, J. S., & Tyree, M. T. (1990). Water-stress-induced xylem embolism in three species of conifers. Plant, Cell & Environment, 13(5), 427–436. https://doi.org/10.1111/j.1365-3040.1990.tb01319.x

Sperry, J S, Donnelly, J. R., & Tyree, M. T. (1988). A method for measuring hydraulic conductivity and embolism in xylem. Plant, Cell and Environment (Vol. 11).

Sperry, Jhon S, & Tyree, M. T. (1988). Mechanism of Water Stress-Induced Xylem Embolism1. Plant Physiol., 88, 581–587.

Tang, K., Fracasso, A., Struik, P. C., Yin, X., & Amaducci, S. (2018). Water-and nitrogen-use efficiencies of hemp (Cannabis sativa L.) based on whole-canopy measurements and modeling. Frontiers in Plant Science, 9. https://doi.org/10.3389/fpls.2018.00951

Tesfamariam, E. H., Annandale, J. G., & Steyn, J. M. (2010). Water stress effects on winter canola growth and yield. Agronomy Journal, 102(2), 658–666. https://doi.org/10.2134/agronj2008.0043

Thomas, D. S., & Eamus, D. (1999). The influence of predawn leaf water potential on stomatal responses to atmospheric water content at constant C(i) and on stem hydraulic conductance and foliar ABA concentrations. Journal of Experimental Botany, 50(331), 243–251. https://doi.org/10.1093/jxb/50.331.243

Tyree, M., & Ewers, F. (1991). The hydraulic architecture of trees and other woody plants. New Phytologist, 119(3), 345–360. https://doi.org/10.1111/J.1469-8137.1991.TB00035.X

Vilagrosa, A., Chirino, E., Peguero-Pina, J.-J., Barigah, S., Cochard, H., & Gil-Pelegrin, E. (2020). Xylem cavitation and embolism in plants living in water-limited ecosystems. https://doi.org/10.1007/978-3-642-32653-0_3ï

Zuardi, A. W. (2005). History of cannabis as a medicine: a review.

